# Genetic structure of invasive baby’s breath (*Gypsophila paniculata*) populations in a freshwater Michigan dune system

**DOI:** 10.1101/401950

**Authors:** Hailee B. Leimbach – Maus, Syndell R. Parks, Charlyn G. Partridge

**Author notes:** Corresponding author: Hailee B. Leimbach – Maus, Annis Water Resources Institute, Grand Valley State University (AWRI-GVSU), 740 W. Shoreline Dr., Muskegon, MI 49441 USA, (no, Email addresses.

## Abstract

Coastal sand dunes are dynamic ecosystems with elevated levels of disturbance, and as such they are highly susceptible to plant invasions. One such invasion that is of major concern to the Great Lakes dune systems is that of perennial baby’s breath (*Gypsophila paniculata*). The invasion of baby’s breath negatively impacts native species such as the federal threatened Pitcher’s thistle (*Cirsium pitcheri*) that occupy the open sand habitat of the Michigan dune system. Our research goals were to (1) quantify the genetic diversity of invasive baby’s breath populations in the Michigan dune system, and (2) estimate the genetic structure of these invasive populations. We analyzed 12 populations at 14 nuclear and 2 chloroplast microsatellite loci. We found strong genetic structure among populations of baby’s breath sampled along Michigan’s dunes (global F_ST_ = 0.228), and also among two geographic regions that are separated by the Leelanau peninsula. Pairwise comparisons using the nSSR data among all 12 populations yielded significant F_ST_ values. Results from a Bayesian clustering analysis suggest two main population clusters. Isolation by distance was found over all 12 populations (R = 0.755, P < 0.001) and when only cluster 2 populations were included (R = 0.523, P = 0.030); populations within cluster 1 revealed no significant relationship (R = 0.205, P = 0.494). Private nSSR alleles and cpSSR haplotypes within each cluster suggest the possibility of at least two separate introduction events to Michigan.

## INTRODUCTION

Coastal sand dunes are dynamic ecosystems. Both the topography and biological community are shaped by disturbance from fluctuations in water levels, weather patterns, and storm events (Arbogast and Loope 1999, Everard et al. 2010, Blumer et al. 2012). In these primary successional systems, vegetation plays an imperative role in trapping sand and soil, both of which accumulate over time and result in sand stabilization and dune formation (Cowles 1899, Olson 1958, Arbogast 2015). Much of the vegetative community native to coastal dune systems is adapted to the harsh conditions posed by the adjacent coast, and some species require early successional, open habitat to thrive (Albert 2000, Everard et al. 2010). For example, dune species such as Marram grass (*Ammophila brevigulata*), Lake Huron tansy (*Tanacetum huronense*), and Pitcher’s thistle (*Cirsium pitcheri*) are adapted to sand burial and will continue to grow above the sand height as it accumulates (Albert 2000). It is the heterogeneous topography and successional processes due to continuous disturbance that makes dune systems so unique (Everard et al.2010).

Because coastal dune ecosystems have naturally elevated levels of disturbance, they are highly susceptible to plant invasions (Jorgensen and Kollman 2009, Carranza et al. 2010, Rand et al. 2015). Invasive plant species are known to be adept at colonizing disturbed areas, and in sparsely-vegetated dune systems that are often in early stages of succession, the opportunities for invasive colonizers are great (Cowles 1899, Grime 1979, Baker 1986, Sakai et al. 2001). Coastal dune systems also typically have a gradient of increasing stages of succession (Cusseddu et al.2016) and this heterogeneous structure can further promote various stages of an invasion, such as colonization, dispersal, and range expansion (With 2002, Theoharides and Dukes 2007). Within the MI dunes system, these successional processes have resulted in a patchwork pattern with alternating areas of open dune habitat, interdunal swales, shrub-scrub, and forested pockets scattered across the landscape (Cowles 1899, Albert 2000, Blumer et al. 2012). This landscape structure can play an important role in shaping species migration, invasive spread, and population demographics (With 2002, Theoharides and Dukes 2007, Jorgensen and Kollman 2009), thus potentially driving patterns of population structure for invasive species. However, management of dune communities can also have a strong impact on invasive populations, as well as the native plant community and the landscape they are invading. Invasive beach grasses *Ammophila breviligulata* and *A. arenaria,* and the management practices used to reduce their impact, led to changes in the morphology of the coastal dune ecosystem by decreasing the maximum dune elevation (Zarnetske et al. 2010). Thus, just as a landscape can shape invasive populations, a plant invasion can also significantly alter the dune landscape (Grime 1979, Cowles 1899, Sakai et al. 2001, Zarnetske et al. 2010).

In addition to the landscape, demographic processes during a species’ invasion also shape the genetic structure observed in contemporary populations. Multiple separate introduction events can result in contemporary populations that are genetically distinct from one another and from the native range (Dlugosch and Parker 2008, Crosby et al. 2014, Hagenblad et al. 2015). Bottleneck events during an introduction can further limit the genetic variation in the invasive range, though this has not necessarily been found to limit the success of an invader (Dlugosch and Parker 2008, Xu et al. 2015). Additionally, genetic admixture and inbreeding can shape the structure of populations, and the effect of these processes can be further influenced by the landscape structure and habitat heterogeneity (Crosby et al. 2014, Nagy and Korpelainen 2014, Moran et al. 2017, Bustamante et al. 2018).

Perennial baby’s breath (*Gypsophila paniculata*) has been identified as a species of concern due to its impact on the integrity of the Michigan dune system (DNR 2015). A perennial iteroparous forb native to the Eurasian steppe region (Darwent and Coupland 1966, Darwent 1975), baby’s breath has been found to negatively impact the coastal dune community in Michigan by crowding out sensitive species such as Pitcher’s thistle (*Cirsium pitcheri*) through direct competition for limited resources, forming monotypic stands in the open dune habitat, preventing the reestablishment of native species, and limiting pollinator visits to native species (Baskett et al. 2011, Jolls et al. 2015, Emery and Doran 2013). Baby’s breath dispersal is thought to be primarily wind-driven (Darwent and Coupland 1966), which is also the mechanism that shapes the dunes. Following seed maturity, the stems of baby’s breath individuals become dry and brittle, breaking at the caudex and forming tumbleweed masses that can disperse roughly 10 000 seeds per plant up to 1 km (Darwent and Coupland 1966, Darwent 1975). Due to the topography and the heterogeneous habitat of the dune systems, the wind patterns of this landscape have the potential to shape the structure of invasive baby’s breath populations. Wind can drive the direction and distance that baby’s breath tumbleweeds disperse, and it is possible that wind patterns could both promote gene flow or limit it by driving tumbleweeds into undesirable habitat. Additionally, the steep topography in parts of the dunes could further prevent tumbleweeds from dispersing significant distances. With these interactive processes in mind, this study explored the genetic structure of invasive populations of baby’s breath within the Michigan coastal dune system. The goals of this research were to (1) quantify the genetic diversity of invasive baby’s breath populations in the Michigan dune system, and (2) estimate the genetic structure of these invasive populations. By estimating the genetic diversity and structure of these invasive populations, we can better understand the impact the dune landscape and its dynamic processes have on this plant invasion.

## METHODS

### Study area and sample collection

To determine the population structure of baby’s breath on a regional scale in Michigan, we collected leaf tissue from plants at 12 different sites in the summers of 2016-2017. All sites were located in areas of known infestation along the dune system of Michigan (Figure 1), and the majority have a history of treatment primarily by The Nature Conservancy, the Grand Traverse Regional Land Conservancy, and the National Park Service (TNC 2013). Eleven sites were located along Lake Michigan in the northwest lower peninsula of Michigan, and one was located on Lake Superior in the upper peninsula. Seven of these sites are located within Sleeping Bear Dunes National Lakeshore (hereafter Sleeping Bear Dunes or SBDNL), which contains one of the largest infestations within the region. We collected leaf tissue samples (5-10 leaves per individual) from a minimum of 20 individuals per site (maximum of 35), and stored them in individual coin envelopes in silica gel until DNA extractions took place (total *n* = 313). Site locations in Michigan (Supplemental A) were separated by a minimum of 10 km and a maximum of 202 km. We subjectively chose individuals to be sampled by identifying a visibly infested area, selecting individuals regardless of size, and walking a minimum of ca. 5 meters in any direction before choosing another plant to minimize the chance of sampling closely related individuals. We observed that the population distributions at the Petoskey State Park and Grand Marais sites were smaller and patchier than the others (ca. 60 individuals total), so we conducted sampling more opportunistically. This opportunistic sampling involved collecting tissue from individuals that were less than 5 m apart, and in some areas sampling from all individuals (ca. 3– 4 individuals) within a small patch (ca. 5m × 5m).

**Figure 1.**
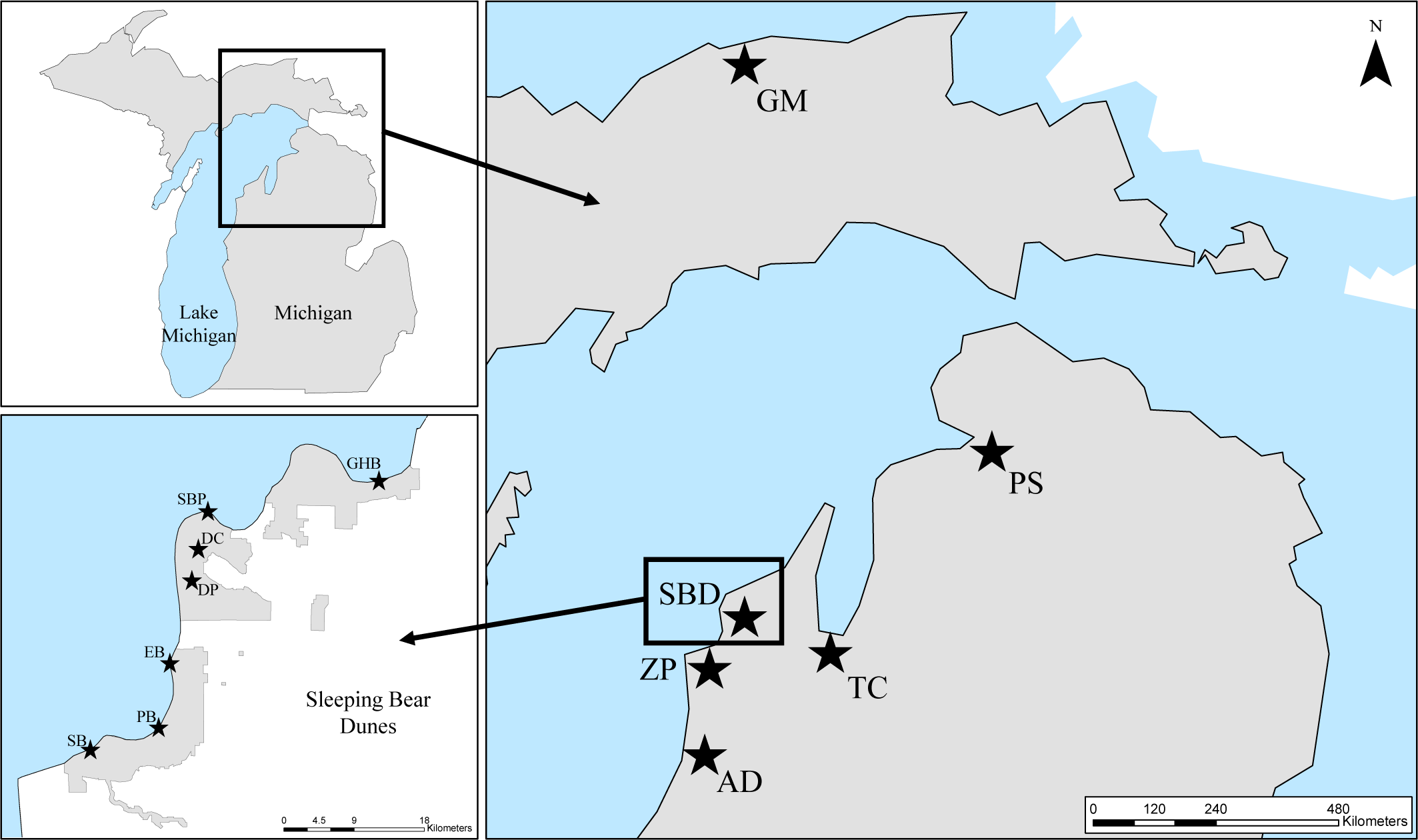
Map of baby’s breath sampling locations in Michigan. Seven were located throughout Sleeping Bear Dunes National Lakeshore. Park boundary delineated by grey shading in bottom left panel. Sampling location codes: Grand Marais (GM), Petoskey State Park (PS), Traverse City (TC), Good Harbor Bay (GHB), Sleeping Bear Point (SBP), Dune Climb (DC), Dune Plateau (DP), Empire Bluffs (EB), Platte Bay (PB), South Boundary (SB), Zetterberg Preserve (ZP), Arcadia Dunes (AD).

### Microsatellite genotyping

We extracted genomic DNA from all samples using DNeasy plant mini kits (QIAGEN, Hilden, Germany) and followed supplier’s instructions with minor modifications, including an extra wash step with AW2 buffer. We then ran the extracted DNA twice through Zymo OneStep PCR Inhibitor Removal Columns (Zymo, Irvine, CA) and quantified the concentrations on a Nanodrop 2000 (Waltham, Massachusetts, USA). We included deionized water controls in each extraction as a quality control for contamination.

We amplified samples at 14 polymorphic nuclear microsatellite loci (hereafter nSSRs) that were developed specifically for *G. paniculata* using Illumina sequencing technology (Table1) (Leimbach-Maus et al. 2018). We conducted polymerase chain reactions (PCR) using a forward primer with a 5’-fluorescent labeled dye (6-FAM, VIC, NED, or PET) (Applied Biosystems, Foster City, CA) and an unlabeled reverse primer. PCR reactions consisted of 1x KCl buffer, 2.0-2.5 mM MgCl_2_ depending on the locus (Table 1), 300 µM dNTP, 0.08 mg/mL BSA, 0.4 µM forward primer fluorescently labeled with either FAM, VIC, NED, or PET, 0.4 µM reverse primer, 0.25 units of Taq polymerase, and a minimum of 50 ng DNA template, all in a 10.0 µL reaction volume. The thermal cycle profile consisted of denaturation at 94°C for 5 minutes followed by 35 cycles of 94°C for 1 minute, annealing at 62°C for 1 min, extension at 72°C for 1 min, and a final elongation step of 72°C for 10 minutes.

**Table 1.**
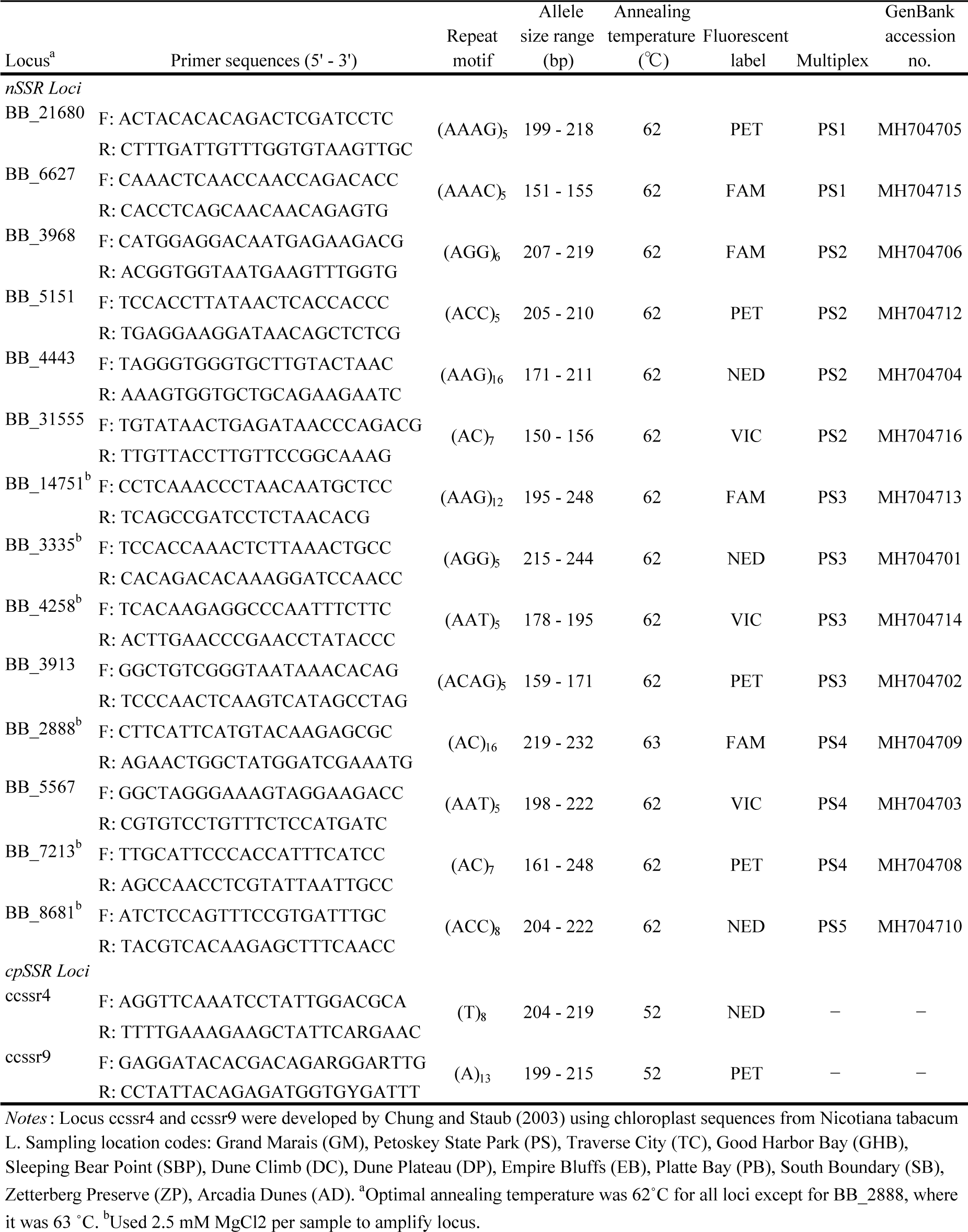
Characteristics of 14 nSSR loci developed for baby’s breath and 2 universal cpSSR loci used in this study.

Each sample was also amplified at 2 universal chloroplast microsatellite loci (hereafter cpSSRs) previously developed for *Nicotiana tabacum* L. (Chung and Staub 2003) (ccssr4,ccssr9) (Table 1). PCR reactions were conducted using a forward primer with a 5’-fluorescent labeled dye and an unlabeled reverse primer. PCR reactions for the cpSSRs are the same as detailed above for the nuclear loci. The thermal cycler profile for cpSSRs is as follows: denaturation at 94°C for 5 minutes followed by 30 cycles of 94°C for 1 minute, annealing at 52°C for 1 minute, extension at 72°C for 1 minute, and a final elongation step of 72°C for 8 minutes (modified from Calistri et al. 2014).

We determined successful amplification by visualizing the amplicons on a 2% agarose gel stained with ethidium bromide. We multiplexed PCR amplicons according to dye color and allele size range (Table 1), added LIZ Genescan 500 size standard, denatured with Hi-Di Formamide at 94°C for four minutes, and then performed fragment analysis on an ABI3130xl Genetic Analyzer (Applied Biosystems) following instrument protocols. We genotyped individuals using the automatic binning procedure on Genemapper v5 (Applied Biosystems), and constructed bins following the Genemapper default settings. To account for the risk of genotyping error when relying on an automated allele-calling procedure, we visually verified that all individuals at all loci were correctly binned to minimize errors caused by stuttering, low heterozygote peak height ratios, and split peaks (DeWoody et al. 2006, Guichoux et al. 2011).

### Quality control

Prior to any analysis, we used multiple approaches to check for scoring errors (DeWoody et al. 2006). We checked nSSR genotypes for null alleles and potential scoring errors due to stuttering and large allele dropout using the software Micro-Checker v2.2.3 (Van Oosterhout et al. 2004, Van Oosterhout et al. 2006). Prior to marker selection, the loci used in this study were previously checked for linkage disequilibrium (Leimbach-Maus et al. 2018). We checked for heterozygote deficiencies in the package STRATAG in the R statistical program. We then screened our data for individuals with more than 20% missing loci and for loci with more than 10% missing individuals (Gomes et al. 1999, Archer et al. 2016). We found none, so all individuals and loci remained for further analyses. In addition, we genotyped 95 individuals twice to ensure consistent allele calls. For the nSSR dataset, we used Genepop 4.2 (Raymond and Rousset 1995, Rousset 2008) to perform an exact test of Hardy-Weinberg Equilibrium (HWE) with 1,000 batches of 1,000 Markov Chain Monte Carlo iterations (Gomes et al. 1999). We also checked for loci out of HWE in more than 60% of the populations; however, there were none.

### nSSR genetic diversity

We calculated the total number of alleles per sampling location, private alleles, observed and expected heterozygosity in GenAlEx 6.502 (Peakall and Smouse 2006, 2012), and estimated the inbreeding coefficient (F_IS_) in Genepop 4.2 (Raymond and Rousset 1995, Rousset 2008). We used the package diverSity in the R statistical program to calculate the allelic richness at each sampling location (Keenan et al. 2013).

### nSSR genetic structure

To test for genetic differentiation between all pairs of sampling locations, we calculated Weir and Cockerham’s (1984) pairwise F_ST_ values for 9,999 permutations in GenAlEx 6.502 (Peakall and Smouse 2006, 2012). In the R statistical program, we corrected the p-values using a false discovery rate (FDR) correction (Benjamini and Hochberg 1995). To test how much of the genetic variance can be explained by within and between population variation, we ran an analysis of molecular variance (AMOVA) for 9,999 permutations in GenAlEx 6.502 (Peakall and Smouse 2006, 2012).

To examine the number of genetic clusters among our sampling locations, we used the Bayesian clustering program STRUCTURE v2.3.2 (Pritchard et al. 2000). Individuals were clustered assuming the admixture model both with and without a priori sampling locations for a burn-in length of 100,000 before 1,000,000 repetitions of MCMC for 10 iterations at each value of *K* (1 – 16). The default settings were used for all other parameters. We identified the most likely value of *K* using the Ln Pr(X|K) from the STRUCTURE output and the ΔK method from Evanno et al. (2005) in CLUMPAK (Kopelman et al. 2015).

To further explore the genetic structure of these populations, we ran a Principal Coordinates Analysis (PCoA) in GenAlEx 6.50, where the analysis was based on an individual pairwise genotypic distance matrix (Peakall et al. 1995, Smouse and Peakall 1999). To find and describe finer genetic structuring of the nSSR dataset, we performed a discriminant analysis of principal components (DAPC) in the R package *adegenet*, which optimizes among-group variance and minimizes within-group variance (Jombart 2008, Jombart et al. 2010). To identify the number of clusters for the analysis, a Bayesian clustering algorithm was run for values of *K* clusters (1 – 16). We retained a *K*-value of 3 based on the resulting Bayesian Information Criterion for each K-value and the results of the previously run PCoA that suggested 3 clusters may exist within the nSSR data. DAPC can be beneficial, as it can limit the number of principal components (PCs) used in the analysis. It has been shown that retaining too many PCs can lead to over-fitting and instability in the membership probabilities returned by the method (Jombart et al. 2010). Therefore, we performed the cross-validation function to identify the optimal number of PCs to retain. Out of 69 total PCs, the cross-validation function suggested we retain 60 PCs (Jombart et al. 2010). We ran the DAPC using the recommended 60 PCs, but also checked if the general patterns remained with fewer PCs used by running the analysis with incrementally less PCs (45 and 30 PCs). All general patterns of the data in the scatterplots remained consistent despite the decreased PCs; therefore, we chose to use the scatterplot based on 30 PCs, as the benefit of the DAPC for our purposes is to show that the main patterns remain, despite minimization of within population variation (Jombart et al. 2010).

To assess the effect of isolation by distance (IBD), we used a paired Mantel test based on a distance matrix of Slatkin’s transformed F_ST_ (D = F_ST_/(1 – F_ST_)) (Slatkin 1995) and a geographic distance matrix for 9,999 permutations in GenAlEx 6.502, and the analysis follows Smouse et al. (1986) and Smouse and Long (1992). The mean geographic center was generated for each sampling location in ArcGIS software (ESRI^TM^ 10.4.1, Redlands, CA), and the latitude and longitude of these points was then used to construct a matrix of straight line distances in km between each sampling location. The reported p-values are based on a one-sided alternative hypothesis (H_1_: R > 0). A Mantel test was run for all sampling locations together, and a test was also run separately for populations within each cluster identified in the STRUCTURE analysis.

### cpSSR genetic diversity

For the cpSSR dataset, we used the program HAPLOTYPE ANALYSIS v1.05 (Eliades and Eliades 2009) to calculate the number of haplotypes, haplotype richness, private haplotypes, and haploid diversity. To visualize patterns in the cpSSR dataset, we created a minimum spanning network in the R package poppr (Kamvar et al. 2014). Nei’s genetic distance was used as the basis for the network with a random seed of 9,999.

### nSSR and cpSSR genetic structure

In order to compare the population structure of the nSSR and cpSSR data, we used the Φ_ST_ distance matrix for both datasets and ran an AMOVA. The population pairwise Φ_ST_ matrix facilitates comparison of molecular variance between codominant and dominant data by suppressing within individual variation, thus allowing for the comparison between varying mutation rates (Weir and Cockerham 1984, Excoffier et al. 1992). To test how much genetic variation could be explained by within populations, between populations, and between regions (genetic clusters identified through STRUCTURE analysis) for both datasets, we ran an AMOVA for 9,999 permutations in GenAlEx 6.502 (Peakall and Smouse 2006, 2012).

## RESULTS

### Microsatellite genotyping and genetic diversity

We genotyped 313 individuals from 12 locations at 14 nSSR loci (Table 1). No loci showed evidence for null alleles across all populations, there were no loci with more than 4 populations significantly out of HWE (less than 30% of populations) (Table 2), and no loci significantly deviated from linkage equilibrium across all populations. The nSSR loci were moderately polymorphic, and the number of alleles per locus per population ranged from 1 – 11, with a total of 85 alleles across 14 loci. Allelic richness (A_R_) ranged from 2.32 – 4.21 per population with a mean of 3.53, and GM, PS, and TC populations exhibited lower levels of A_R_ than the other populations. Of the 6 private nSSR alleles identified, 5 were at low frequencies – occurring in five or fewer individuals, but the private nSSR allele in the GM population occurred in over 60% of individuals. Overall, the observed heterozygosity (H_O_) averaged over loci for each population ranged from 0.25 – 0.56 with a mean of 0.46, and the 3 northernmost populations (GM, PS, TC) had lower diversity in general. Expected heterozygosity (H_E_) ranged from 0.30 – 0.57 across populations, with a mean of 0.49. GM and AD populations deviated significantly from HWE (P < 0.05). GM had a higher inbreeding coefficient (Table 2), but this could be attributed to our limited area in which to sample.

**Table 2.** Genetic diversity indices for baby’s breath from each sampling location at 14 nSSRs and 2 cpSSRs.

|  | Sampling Locations |  |  |  |  |  |  |  |  |  |  |  |
| --- | --- | --- | --- | --- | --- | --- | --- | --- | --- | --- | --- | --- |
|  | GM | PS | TC | GHB | SBP | DC | DP | EB | PB | SB | ZP | AD |
| **nSSR Loci** |  |  |  |  |  |  |  |  |  |  |  |  |
| *BB_21680* |  |  |  |  |  |  |  |  |  |  |  |  |
| *N* | 35 | 30 | 30 | 20 | 25 | 23 | 30 | 19 | 20 | 20 | 30 | 30 |
| *N<sub>A</sub>* | 3 | 3 | 4 | 3 | 3 | 4 | 3 | 4 | 3 | 4 | 4 | 3 |
| *H<sub>O</sub>* | 0.286 | 0.667 | 0.300 | 0.450 | 0.440 | 0.522 | 0.500 | 0.421 | 0.500 | 0.400 | 0.500 | 0.700 |
| *H<sub>E</sub>* | 0.411 | 0.539 | 0.534 | 0.540 | 0.582 | 0.481 | 0.540 | 0.676 | 0.524 | 0.510 | 0.548 | 0.546 |
| *F<sub>IS</sub>* | **0.3186** | -0.2198 | **0.4517** | 0.1915 | 0.2636 | -0.0624 | 0.0909 | **0.4000** | 0.0709 | **0.2400** | 0.1040 | -0.2661 |
| *BB_6627* |  |  |  |  |  |  |  |  |  |  |  |  |
| *N* | 35 | 30 | 30 | 20 | 24 | 23 | 30 | 20 | 20 | 20 | 30 | 30 |
| *N<sub>A</sub>* | 2 | 1 | 2 | 2 | 2 | 2 | 2 | 2 | 2 | 2 | 2 | 2 |
| *H<sub>O</sub>* | 0.086 | 0.000 | 0.167 | 0.500 | 0.333 | 0.435 | 0.500 | 0.250 | 0.450 | 0.550 | 0.300 | 0.467 |
| *H<sub>E</sub>* | 0.082 | 0.000 | 0.153 | 0.480 | 0.330 | 0.499 | 0.495 | 0.219 | 0.489 | 0.439 | 0.255 | 0.464 |
| *F<sub>IS</sub>* | -0.0303 | - | -0.0741 | -0.0160 | 0.0108 | 0.1506 | 0.0068 | -0.1176 | 0.1047 | -0.2294 | -0.1600 | 0.0122 |
| *BB_3968* |  |  |  |  |  |  |  |  |  |  |  |  |
| *N* | 35 | 30 | 30 | 20 | 25 | 23 | 30 | 20 | 20 | 20 | 30 | 30 |
| *N<sub>A</sub>* | 3 | 2 | 1 | 4 | 3 | 3 | 4 | 4 | 3 | 4 | 3 | 2 |
| *H<sub>O</sub>* | 0.143 | 0.067 | 0.000 | 0.400 | 0.240 | 0.304 | 0.367 | 0.450 | 0.550 | 0.400 | 0.500 | 0.133 |
| *H<sub>E</sub>* | 0.207 | 0.064 | 0.000 | 0.476 | 0.246 | 0.334 | 0.414 | 0.475 | 0.509 | 0.345 | 0.418 | 0.180 |
| *F<sub>IS</sub>* | **0.3241** | -0.0175 | - | 0.1850 | 0.0432 | 0.1098 | 0.1320 | 0.0782 | -0.0556 | -0.1343 | -0.1805 | 0.2750 |
| *BB_5151* |  |  |  |  |  |  |  |  |  |  |  |  |
| *N* | 35 | 29 | 30 | 19 | 25 | 23 | 30 | 19 | 20 | 20 | 30 | 28 |
| *N<sub>A</sub>* | 2 | 2 | 2 | 2 | 2 | 2 | 2 | 2 | 2 | 2 | 2 | 2 |
| *H<sub>O</sub>* | 0.057 | 0.034 | 0.133 | 0.158 | 0.200 | 0.391 | 0.467 | 0.526 | 0.400 | 0.400 | 0.200 | 0.179 |
| *H<sub>E</sub>* | 0.056 | 0.034 | 0.231 | 0.494 | 0.449 | 0.466 | 0.491 | 0.465 | 0.480 | 0.375 | 0.180 | 0.499 |
| *F<sub>IS</sub>* | -0.0149 | -0.018 | 0.4369 | **0.6949** | **0.5683** | 0.1818 | 0.0667 | -0.1043 | 0.1915 | -0.0411 | -0.0943 | **0.6530** |
| *BB_4443* |  |  |  |  |  |  |  |  |  |  |  |  |
| *N* | 35 | 30 | 30 | 20 | 25 | 23 | 30 | 19 | 20 | 19 | 30 | 30 |
| *N<sub>A</sub>* | 3 | 5 | 3 | 9 | 9 | 9<sup>A</sup> | 9 | 5 | 11 | 7 | 10<sup>A</sup> | 5 |
| *H<sub>O</sub>* | 0.257 | 0.800 | 0.400 | 0.800 | 0.640 | 0.783 | 0.767 | 0.842 | 0.900 | 0.526 | 0.667 | 0.567 |
| *H<sub>E</sub>* | 0.338 | 0.701 | 0.399 | 0.808 | 0.769 | 0.778 | 0.758 | 0.749 | 0.853 | 0.593 | 0.651 | 0.663 |
| *F<sub>IS</sub>* | **0.2537** | -0.1244 | 0.0156 | 0.0349 | 0.1804 | 0.0161 | 0.0052 | -0.0971 | -0.0301 | 0.2193 | -0.0078 | 0.1623 |
| *BB_31555* |  |  |  |  |  |  |  |  |  |  |  |  |
| *N* | 28 | 30 | 30 | 20 | 25 | 23 | 30 | 20 | 20 | 20 | 30 | 30 |
| *N<sub>A</sub>* | 1 | 2 | 2 | 3 | 4 | 4 | 4 | 4 | 4 | 4 | 4 | 3 |
| *H<sub>O</sub>* | 0.000 | 0.233 | 0.333 | 0.650 | 0.800 | 0.478 | 0.600 | 0.650 | 0.750 | 0.500 | 0.600 | 0.467 |
| *H<sub>E</sub>* | 0.000 | 0.255 | 0.278 | 0.609 | 0.727 | 0.650 | 0.614 | 0.654 | 0.745 | 0.583 | 0.663 | 0.545 |
| *F<sub>IS</sub>* | - | 0.1018 | -0.1837 | -0.0422 | -0.0799 | 0.2851 | 0.0396 | 0.0314 | 0.0189 | 0.1667 | 0.1122 | 0.1603 |
| *BB_14751* |  |  |  |  |  |  |  |  |  |  |  |  |
| *N* | 34 | 30 | 30 | 20 | 25 | 23 | 30 | 20 | 20 | 20 | 28 | 30 |
| *N<sub>A</sub>* | 5<sup>A</sup> | 3 | 4 | 7 | 6 | 7 | 8 | 9 | 9 | 10 | 6 | 7 |
| *H<sub>O</sub>* | 0.676 | 0.200 | 0.467 | 0.650 | 0.600 | 0.478 | 0.633 | 0.800 | 0.600 | 0.650 | 0.750 | 0.467 |
| *H<sub>E</sub>* | 0.666 | 0.287 | 0.548 | 0.714 | 0.633 | 0.618 | 0.769 | 0.790 | 0.621 | 0.646 | 0.675 | 0.762 |
| *F<sub>IS</sub>* | **-0.0013** | 0.3176 | -0.0899 | 0.1099 | 0.0722 | 0.2473 | 0.1933 | 0.0130 | 0.0579 | 0.0159 | -0.0925 | 0.2632 |
| *BB_3335* |  |  |  |  |  |  |  |  |  |  |  |  |
| *N* | 33 | 30 | 30 | 20 | 25 | 23 | 30 | 20 | 20 | 20 | 30 | 30 |
| *N<sub>A</sub>* | 5 | 3 | 4 | 8 | 8 | 9 | 7 | 5 | 9 | 10 | 8 | 6 |
| *H<sub>O</sub>* | 0.242 | 0.500 | 0.433 | 0.800 | 0.760 | 0.826 | 0.667 | 0.800 | 0.750 | 0.850 | 0.767 | 0.600 |
| *H<sub>E</sub>* | 0.403 | 0.562 | 0.369 | 0.818 | 0.732 | 0.827 | 0.817 | 0.694 | 0.808 | 0.815 | 0.789 | 0.709 |
| *F<sub>IS</sub>* | **0.4115** | **0.1265** | -0.1564 | 0.0470 | -0.0179 | 0.0234 | **0.2000** | -0.1280 | 0.0967 | -0.0173 | 0.0451 | 0.1701 |

**Table 2 (Cont)** Genetic diversity indices for baby's breath from each sampling location at 14 nSSRs and 2 cpSSRs.
|  | Sampling Locations |  |  |  |  |  |  |  |  |  |  |  |
| --- | --- | --- | --- | --- | --- | --- | --- | --- | --- | --- | --- | --- |
|  | GM | PS | TC | GHB | SBP | DC | DP | EB | PB | SB | ZP | AD |
| **nSSR Loci** |  |  |  |  |  |  |  |  |  |  |  |  |
| *BB_4258* |  |  |  |  |  |  |  |  |  |  |  |  |
| *N* | 35 | 30 | 30 | 20 | 25 | 23 | 30 | 20 | 20 | 20 | 30 | 30 |
| *N<sub>A</sub>* | 1 | 1 | 2 | 3<sup>A</sup> | 2 | 2 | 2 | 1 | 2 | 4<sup>A</sup> | 3 | 2 |
| *H<sub>O</sub>* | 0.000 | 0.000 | 0.133 | 0.150 | 0.240 | 0.087 | 0.033 | 0.000 | 0.150 | 0.250 | 0.400 | 0.300 |
| *H<sub>E</sub>* | 0.000 | 0.000 | 0.124 | 0.141 | 0.269 | 0.083 | 0.033 | 0.000 | 0.139 | 0.228 | 0.326 | 0.339 |
| *F<sub>IS</sub>* | – | – | –0.0545 | –0.0364 | 0.1273 | –0.0233 | –0.017 | – | –0.0556 | –0.0734 | –0.2104 | 0.1329 |
| *BB_3913* |  |  |  |  |  |  |  |  |  |  |  |  |
| *N* | 35 | 30 | 30 | 20 | 25 | 23 | 30 | 20 | 20 | 20 | 30 | 30 |
| *N<sub>A</sub>* | 3 | 2 | 3 | 4 | 4 | 4 | 4 | 4 | 4 | 4 | 4 | 2 |
| *H<sub>O</sub>* | 0.171 | 0.133 | 0.300 | 0.400 | 0.480 | 0.565 | 0.667 | 0.550 | 0.500 | 0.550 | 0.600 | 0.467 |
| *H<sub>E</sub>* | 0.207 | 0.124 | 0.292 | 0.471 | 0.537 | 0.619 | 0.578 | 0.614 | 0.494 | 0.621 | 0.638 | 0.444 |
| *F<sub>IS</sub>* | 0.1873 | –0.0545 | –0.0166 | 0.1762 | 0.1259 | 0.1090 | –0.1373 | **0.1292** | 0.0130 | 0.1399 | 0.0769 | –0.0331 |
| *BB_2888* |  |  |  |  |  |  |  |  |  |  |  |  |
| *N* | 35 | 29 | 30 | 20 | 25 | 23 | 30 | 20 | 20 | 20 | 30 | 30 |
| *N<sub>A</sub>* | 4 | 2 | 3 | 4 | 6 | 6 | 5 | 6 | 5 | 6<sup>A</sup> | 6 | 5 |
| *H<sub>O</sub>* | 0.657 | 0.586 | 0.600 | 0.800 | 0.680 | 0.913 | 0.833 | 0.750 | 0.800 | 0.800 | 0.900 | 0.667 |
| *H<sub>E</sub>* | 0.594 | 0.498 | 0.651 | 0.734 | 0.724 | 0.772 | 0.793 | 0.768 | 0.746 | 0.734 | 0.786 | 0.589 |
| *F<sub>IS</sub>* | –0.0922 | –0.1610 | 0.0953 | –0.0648 | 0.0811 | –0.1608 | –0.0335 | 0.0484 | –0.0465 | –0.0648 | –0.1282 | –0.1154 |
| *BB_5567* |  |  |  |  |  |  |  |  |  |  |  |  |
| *N* | 35 | 30 | 30 | 20 | 25 | 23 | 30 | 20 | 20 | 20 | 30 | 30 |
| *N<sub>A</sub>* | 4 | 3 | 4 | 5 | 3 | 3 | 4 | 3 | 4 | 3 | 5 | 5 |
| *H<sub>O</sub>* | 0.629 | 0.533 | 0.567 | 0.700 | 0.480 | 0.609 | 0.667 | 0.400 | 0.550 | 0.400 | 0.600 | 0.767 |
| *H<sub>E</sub>* | 0.716 | 0.613 | 0.562 | 0.745 | 0.614 | 0.474 | 0.604 | 0.374 | 0.589 | 0.371 | 0.613 | 0.716 |
| *F<sub>IS</sub>* | **0.1368** | 0.1463 | 0.0080 | 0.0859 | 0.2371 | –0.2649 | **–0.0872** | –0.0447 | 0.0913 | –0.0519 | 0.0387 | –0.0545 |
| *BB_7213* |  |  |  |  |  |  |  |  |  |  |  |  |
| *N* | 35 | 30 | 30 | 20 | 25 | 23 | 30 | 20 | 20 | 20 | 30 | 30 |
| *N<sub>A</sub>* | 3 | 2 | 3 | 3 | 4 | 3 | 3 | 3 | 3 | 5 | 5 | 3 |
| *H<sub>O</sub>* | 0.229 | 0.367 | 0.100 | 0.500 | 0.640 | 0.391 | 0.500 | 0.400 | 0.650 | 0.500 | 0.633 | 0.667 |
| *H<sub>E</sub>* | 0.359 | 0.375 | 0.096 | 0.434 | 0.642 | 0.638 | 0.565 | 0.386 | 0.611 | 0.499 | 0.599 | 0.633 |
| *F<sub>IS</sub>* | **0.3754** | 0.0392 | –0.0235 | –0.1276 | 0.0241 | **0.4054** | 0.1317 | –0.0100 | –0.0378 | 0.0231 | –0.0406 | –0.0366 |
| *BB_8681* |  |  |  |  |  |  |  |  |  |  |  |  |
| *N* | 35 | 28 | 30 | 20 | 24 | 22 | 30 | 19 | 20 | 20 | 30 | 30 |
| *N<sub>A</sub>* | 3 | 3 | 2 | 3 | 3 | 3 | 4 | 2 | 3 | 3 | 4 | 3 |
| *H<sub>O</sub>* | 0.114 | 0.357 | 0.333 | 0.500 | 0.500 | 0.136 | 0.400 | 0.211 | 0.250 | 0.300 | 0.467 | 0.600 |
| *H<sub>E</sub>* | 0.109 | 0.523 | 0.320 | 0.395 | 0.469 | 0.206 | 0.456 | 0.188 | 0.265 | 0.464 | 0.502 | 0.438 |
| *F<sub>IS</sub>* | –0.0342 | **0.3333** | –0.0247 | –0.2418 | –0.0455 | 0.3571 | 0.1397 | –0.0909 | 0.0821 | **0.3753** | 0.0866 | –0.3558 |
| *A<sub>R</sub> across loci* | 2.660 | 2.320 | 2.540 | 3.970 | 3.750 | 3.920 | 3.990 | 3.660 | 4.070 | 4.210 | 4.190 | 3.120 |
| **cpSSR Loci** |  |  |  |  |  |  |  |  |  |  |  |  |
| *ccssr4* |  |  |  |  |  |  |  |  |  |  |  |  |
| *N* | 35 | 30 | 29 | 20 | 25 | 23 | 30 | 20 | 20 | 20 | 30 | 30 |
| *N<sub>A</sub>* | 2 | 1 | 1 | 1 | 1 | 1 | 1 | 1 | 1 | 2 | 2 | 1 |
| *H* | 0.25 | 0.00 | 0.00 | 0.00 | 0.00 | 0.00 | 0.00 | 0.00 | 0.00 | 0.32 | 0.50 | 0.00 |
| *ccssr9* |  |  |  |  |  |  |  |  |  |  |  |  |
| *N* | 35 | 30 | 29 | 20 | 25 | 23 | 30 | 20 | 20 | 20 | 30 | 30 |
| *N<sub>A</sub>* | 2<sup>A</sup> | 1 | 1 | 1 | 1 | 1 | 1 | 1 | 1 | 2 | 2 | 1 |
| *H* | 0.25 | 0.00 | 0.00 | 0.00 | 0.00 | 0.00 | 0.00 | 0.00 | 0.00 | 0.10 | 0.12 | 0.00 |
| *N<sub>H</sub> across loci* | 2<sup>P</sup> | 1 | 1 | 1 | 1 | 1 | 1 | 1 | 1 | 3 | 3 | 1 |
| *H<sub>R</sub> across loci* | 0.991 | 0 | 0 | 0 | 0 | 0 | 0 | 0 | 0 | 2 | 1.897 | 0 |

Notes : *N* number of individuals, *N<sub>A</sub>* number of alleles per locus, *H<sub>O</sub>* observed heterozygosity, *H<sub>E</sub>* expected heterozygosity, *F<sub>IS</sub>* inbreeding coefficient (Weir and Cockerham 1984), *A<sub>R</sub>* allelic richness for each population averaged across loci, *H* haploid diversity, *N<sub>H</sub>* number of haplotypes for each population averaged across loci, *H<sub>R</sub>* haplotype richness for each population averaged across loci. Bold values indicate loci that deviated from Hardy-Weinberg equilibrium. <sup>A</sup> denotes a private allele, <sup>P</sup> denotes a private haplotype. Sampling location codes: Grand Marais (GM), Petoskey State Park (PS), Traverse City (TC), Good Harbor Bay (GHB), Sleeping Bear Point (SBP), Dune Climb (DC), Dune Plateau (DP), Empire Bluffs (EB), Platte Bay (PB), South Boundary (SB), Zetterberg Preserve (ZP), Arcadia Dunes (AD).

Both cpSSR loci were polymorphic, with 3 alleles per locus for a total of 6 alleles, and the number of alleles per population ranged from 2 – 4 with an average of 2.50 (Table 2). All alleles together resulted in 5 haplotypes. There were between 1 – 3 haplotypes per population for a haplotype richness ranging from 0.00 – 2.00, with a mean of 0.41 per population. Haploid diversity ranged from 0.00 – 0.58 with a mean of 0.10 per population. One allele and haplotype 2 were both unique to the SB and ZP sampling locations, and another allele and haplotype 4 were both private to five individuals sampled in GM, which occurred in a separate sampling location from the rest of the individuals in GM (Figure 5).

### Genetic structure

The nSSR data suggested that there is strong genetic structure among the populations and regions of baby’s breath sampled along the dunes of western and northern Michigan (global F_ST_ = 0.228). Pairwise comparisons using the nSSR data among all 12 populations yielded significant F_ST_ values after a FDR correction (Benjamini and Hochberg 1995) (Table 3). However, all pairwise comparisons of populations within Sleeping Bear Dunes (GHB, SBP, DC, DP, EB, PB, SB) and nearby ZP displayed relatively lower pairwise F_ST_ values (Table 3), suggesting that there is some gene flow among these populations. The AMOVA based on the nSSR data also found that a significant amount of the genetic variation could be explained by differences between populations in the northern region (GM, PS, TC) and populations in the southern region (GHB,SBP, DC, DP, EB, PB, SB, ZP, AD) (F_CT_ = 0.144, P < 0.0001), as well as among populations within regions (F_SC_ = 0.097, P < 0.0001). However, the majority of the genetic variance was explained by among population differences (F_ST_ = 0.228, P < 0.0001).

**Table 3.** Pairwise F_ST_ values for nSSR data among all sampling locations based on Weir and Cockerham’s (1984) estimate. Darker color – increasing F_ST_ value, lighter color – decreasing F_ST_ value.

|  | GM | PS | TC | GHB | SBP | DC | DP | EB | PB | SB | ZP | AD |
| --- | --- | --- | --- | --- | --- | --- | --- | --- | --- | --- | --- | --- |
| GM | – | – | – | – | – | – | – | – | – | – | – | – |
| PS | 0.221 | – | – | – | – | – | – | – | – | – | – | – |
| TC | 0.147 | 0.121 | – | – | – | – | – | – | – | – | – | – |
| GHB | 0.264 | 0.245 | 0.221 | – | – | – | – | – | – | – | – | – |
| SBP | 0.261 | 0.246 | 0.230 | 0.047 | – | – | – | – | – | – | – | – |
| DC | 0.321 | 0.320 | 0.277 | 0.063 | 0.041 | – | – | – | – | – | – | – |
| DP | 0.272 | 0.265 | 0.246 | 0.026 | 0.029 | 0.025 | – | – | – | – | – | – |
| EB | 0.220 | 0.240 | 0.175 | 0.081 | 0.069 | 0.093 | 0.083 | – | – | – | – | – |
| PB | 0.290 | 0.283 | 0.260 | 0.062 | 0.040 | 0.050 | 0.050 | 0.106 | – | – | – | – |
| SB | 0.253 | 0.266 | 0.207 | 0.094 | 0.057 | 0.077 | 0.069 | 0.068 | 0.066 | – | – | – |
| ZP | 0.240 | 0.238 | 0.211 | 0.071 | 0.040 | 0.090 | 0.055 | 0.071 | 0.082 | 0.017 | – | – |
| AD | 0.298 | 0.254 | 0.231 | 0.121 | 0.121 | 0.137 | 0.123 | 0.170 | 0.128 | 0.132 | 0.128 | – |

*Notes* : All values significant at $P \leq 0.005$ after FDR correction (Benjamini and Hochberg 1995). Sampling location codes: Grand Marais (GM), Petoskey State Park (PS), Traverse City (TC), Good Harbor Bay (GHB), Sleeping Bear Point (SBP), Dune Climb (DC), Dune Plateau (DP), Empire Bluffs (EB), Platte Bay (PB), South Boundary (SB), Zetterberg Preserve (ZP), Arcadia Dunes (AD).

The Bayesian clustering analysis from the program STRUCTURE (Pritchard et al. 2000) partitioned the population into two clusters (*K* = 2) (Figure 2), inferred from both Ln Pr(X|*K*) and Evanno’s Δ*K* (Supplemental B). This analysis was run without inferring any prior information on sampling location, and then again with sampling information as prior. No differences were observed between the two results (without priors shown in Figure 2). Cluster 1 is comprised of the northernmost populations (GM, PS, TC), and cluster 2 includes all other populations.

**Figure 2.**
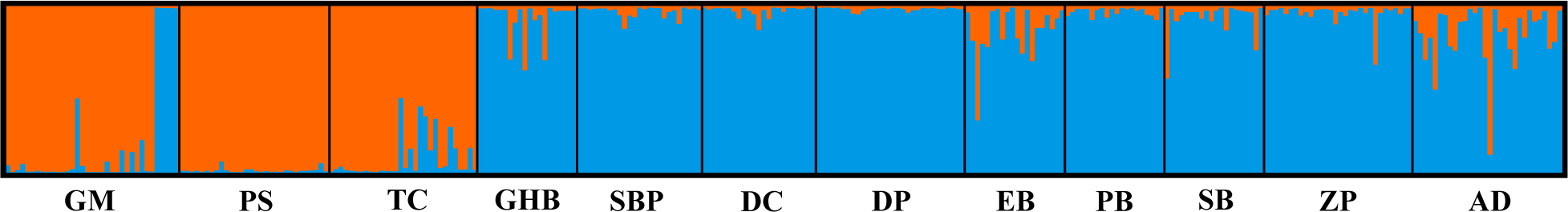
Results from Bayesian cluster analysis based on nSSR data using the program STRUCTURE (Pritchard et al. 2000) indicate (*K* = 2) population clusters of baby’s breath (Pritchard et al. 2000, Evanno et al. 2005). Cluster 1 (left) includes the three northernmost populations, and Cluster 2 (right) includes all other populations. Each individual (*N* = 313) is represented by a line in the plot, and individuals are grouped by population. Sampling location codes: Grand Marais (GM), Petoskey State Park (PS), Traverse City (TC), Good Harbor Bay (GHB), Sleeping Bear Point (SBP), Dune Climb (DC), Dune Plateau (DP), Empire Bluffs (EB), Platte Bay (PB), South Boundary (SB), Zetterberg Preserve (ZP), Arcadia Dunes (AD).

However, five individuals in the GM population (cluster 1) were assigned to cluster 2 (assignment probability > 90%), and these individuals were located at a separate sampling location from the rest in GM. In addition, though there is little admixture overall, several individuals in the GM, TC, EB, and AD populations showed a higher proportion of admixture among the two clusters.

The Principal Coordinates analysis (PCoA) based on an individual pairwise genotypic distance matrix highlighted population substructuring (Supplemental C). Individuals in the AD population expanded along both principal coordinates away from individuals assigned to the original STRUCTURE cluster 2 (Figure 2). In addition, the scatterplot supported the strong grouping of individuals in GM, PS, and TC together.

A Discriminant Analysis of Principal Components (DAPC) scatterplot (Figure 3a) grouped individuals into three clusters along two axes, supporting the substructuring illustrated in the PCoA. While the PCoA illustrated global diversity found in the nSSR dataset, the DAPC optimizes between group variance. Figure 3b shows the overlap between the distributions of individuals in DAPC clusters 2 and 3 along the first discriminant function, suggesting little distance between them. The membership of individuals of each population to the three illustrated clusters can be seen in Figure 3c. This visualization of the data further highlights the more subtle structure of baby’s breath populations in the dunes system of Michigan.

**Figure 3.**
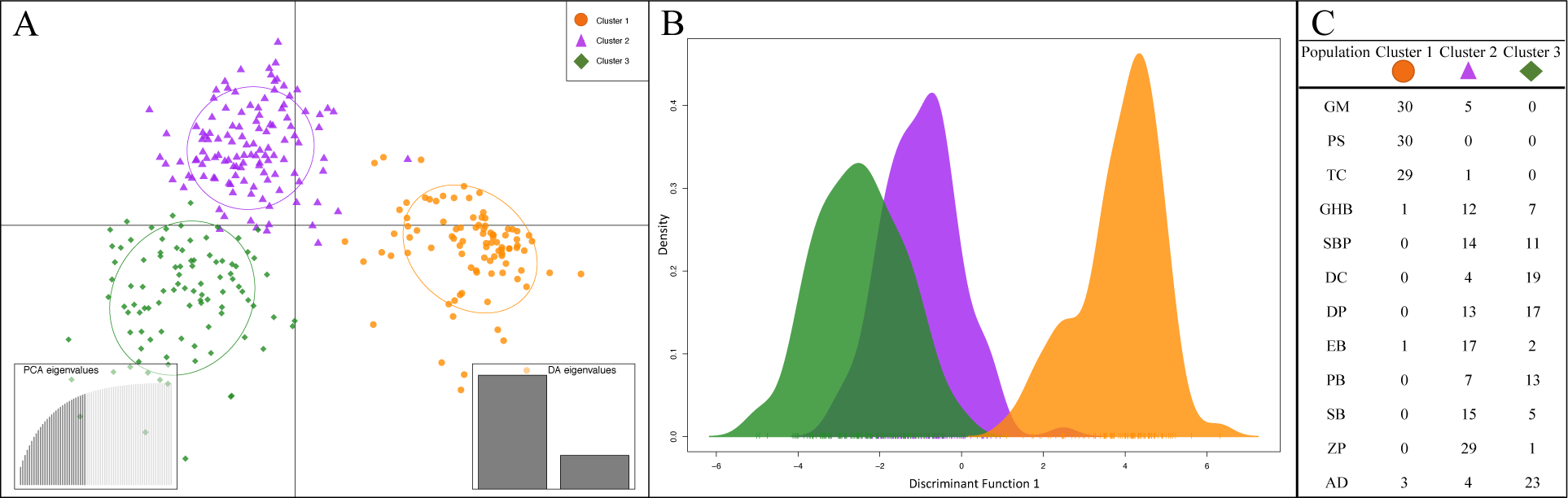
Discriminant analysis of principal components (DAPC) based on baby’s breath nSSR data and calculated in the *adegenet* v2.1.0 (Jombart 2008, Jombart et al. 2010) package for R. (A) Scatterplot of both discriminant function axes; all individuals (*n* = 313) are included and represented by a dot. (B) Plot of DAPC sample distribution on discriminant function 1. (C) Table of individual membership to each DAPC cluster, explained by the PCA eigenvalues used in the DAPC, based on all 69 identified principal components. Sampling location codes: Grand Marais (GM), Petoskey State Park (PS), Traverse City (TC), Good Harbor Bay (GHB), Sleeping Bear Point (SBP), Dune Climb (DC), Dune Plateau (DP), Empire Bluffs (EB), Platte Bay (PB), South Boundary (SB), Zetterberg Preserve (ZP), Arcadia Dunes (AD).

A Mantel test for isolation by distance (IBD) performed over all populations found a significant correlation between genetic and geographic distances (R = 0.755, P < 0.001) (Figure 4a). Upon further exploration of this correlation through separate Mantel tests within each identified STRUCTURE cluster, we found a significant correlation within cluster 2 (Figure 4c) (R = 0.523, P = 0.030), but no significant correlation within cluster 1 (Figure 4b) (R = 0.205, P =0.494).

**Figure 4.**
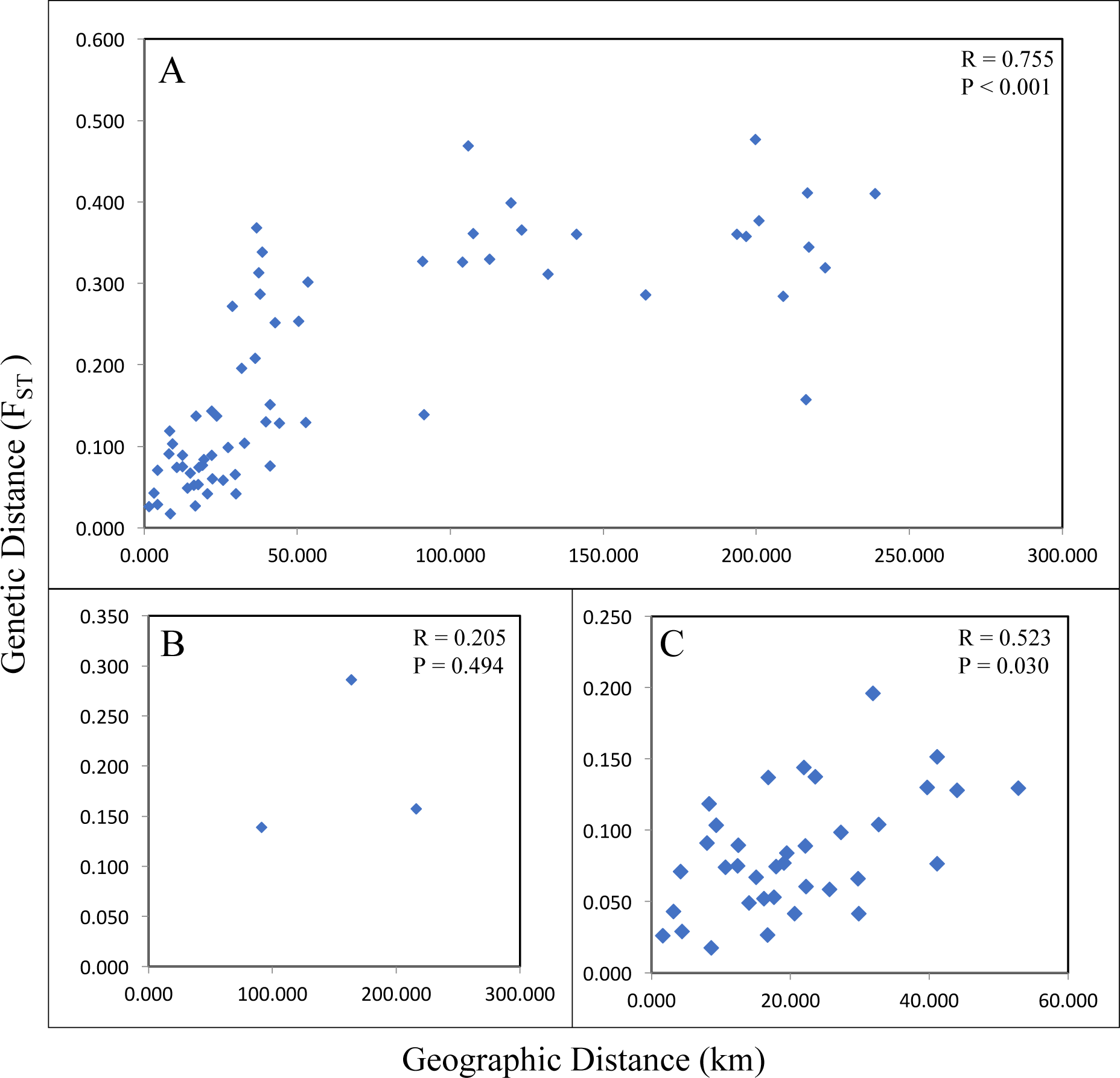
Mantel tests using transformed pairwise population F_ST_ values of nSSR data (Slatkin 1995) and straight-line distances (km) between populations based on the mean center latitude and longitude of each location. (A) Between all populations, (B) between populations in the Northern region (cluster 1), and (C) between populations in the Southern region (cluster 2) identified from the Bayesian clustering analysis. Reported p-values based on the one-sided alternative hypothesis (H^1^: R > 0).

The AMOVA based on **Φ**_ST_ distance (Supplemental D) facilitated the comparison between the nSSR and cpSSR data, which resulted in a significant amount of the genetic variation explained by differences among regions (nSSR **Φ**_CT_ = 0.226, cpSSR **Φ**_CT_ = 0.263), among populations within regions (nSSR **Φ**_SC_ = 0.167, cpSSR **Φ**_SC_ =0.643), and within populations (nSSR **Φ**_ST_ = 0.355, cpSSR **Φ**_ST_ = 0.736) for both data sets (P < 0.0001).

For the cpSSR markers, the minimum spanning network illustrates the distribution of haplotypes across the 12 populations (Figure 5). Five haplotypes were found; Haplotype 1 was the most common, but only occurred in the SBDNL and ZP populations (GHB, SBP, DC, DP, EB, PB, SB, ZP). Haplotype 2 was private to the SB and ZP populations, but rare, occurring in one and two individuals, respective to the populations. Haplotype 3 was private to the five GM individuals located separately from the majority of the other individuals from the GM population. Haplotype 4 was private to SB, ZP, and AD populations, and occurred in all AD individuals, but was less common in the SB and ZP populations. Haplotype 5 was private to GM, PS, and TC populations, occurring in all individuals.

**Figure 5.**
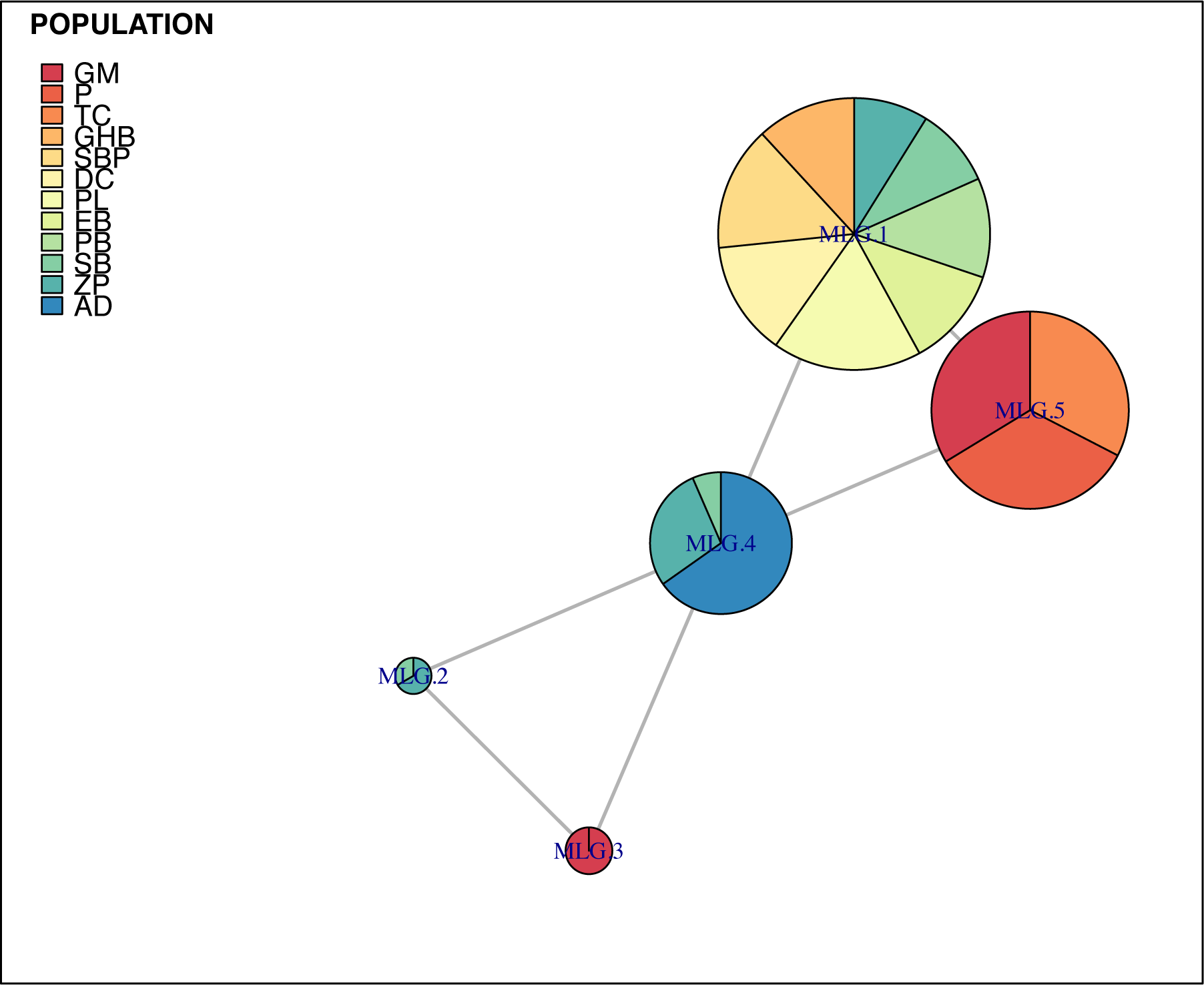
Minimum spanning network based on Nei’s genetic distance (Nei 1972) matrix of baby’s breath cpSSR data. Created in the *poppr* v2.8.0 package (Kamvar et al. 2014) for R. Illustrates the distribution of haplotypes across the 12 populations. Haplotype size indicates frequency in populations. Sampling location codes: Grand Marais (GM), Petoskey State Park (PS), Traverse City (TC), Good Harbor Bay (GHB), Sleeping Bear Point (SBP), Dune Climb (DC), Dune Plateau (DP), Empire Bluffs (EB), Platte Bay (PB), South Boundary (SB), Zetterberg Preserve (ZP), Arcadia Dunes (AD).

## DISCUSSION

The natural disturbance regime of dynamic sand dune systems can result in a pattern of fragmented habitat and often sparse vegetative cover, making dune ecosystems highly susceptible to plant invasions (Jorgensen and Kollman 2009, Carranza et al. 2010, Rand et al. 2015). The topography, geographic distribution of preferred habitat, and disturbance regime in an ecosystem can influence various stages of a species invasion, including where the plant establishes, its dispersal patterns, and how densely it grows (With 2002, Theoharides and Dukes 2007). In addition, the demographic processes of an introduction event can shape contemporary population dynamics (Dlugosch and Parker 2008, Estoup and Guillemaud 2010). The invasion of baby’s breath in the Michigan dune system is an opportunity to better understand the genetic structure of invasive species in this system and how the dynamic landscape of these dunes may be shaping it. Our results indicate variation in genetic diversity among populations, as well as strong genetic structure that clusters individuals into two distinct groups. These two groups are separated by a peninsula that could be limiting gene flow between the two groups, causing this genetic separation.

We observed moderate levels of nuclear and chloroplast genetic diversity across populations of baby’s breath throughout the dune system of Michigan (Table 1). However, genetic diversity in our northern-most populations (Grand Marais, Petoskey State Park, and Traverse City) was typically lower compared to that found in the populations in Sleeping Bear Dunes, Zetterberg Preserve, and Arcadia Dunes. Differences in the level of genetic diversity among these regions could be due to differences in population size. Sleeping Bear Dunes is a largescale infestation and has some of the highest densities of baby’s breath found within the Michigan coastal dunes (TNC 2013), consisting of up to 80% of the vegetation and covering hundreds of acres in some areas. The Grand Marais, Petoskey State Park, and Traverse City populations are much smaller than those found in Sleeping Bear Dunes, with continuous populations often limited to less than 45 acres (TNC 2012 internal report). These smaller populations could be more affected by the impact of genetic drift and potential inbreeding, resulting in the observed lower levels of genetic diversity (Ellstrand and Elam 1993, Young et al. 1996, Keller and Waller 2002).

The level of isolation between Grand Marais, Petoskey State Park, and Traverse City could also be contributing to the lower levels of genetic diversity observed in these areas compared to other populations. F_ST_ values among these three populations ranged from 0.121 – 0.221, which is much higher than the F_ST_ range observed between the Sleeping Bear Dunes, Zetterberg Preserve, and Arcadia Dunes populations (0.041 – 0.137). This suggests that our northern-most populations may have less gene flow between neighboring populations. This could be the result of larger geographic distances between these locations. For example, Grand Marais is located in Michigan’s upper peninsula while Petoskey State Park and Traverse City are located in the lower peninsula. Higher levels of isolation could also be a result of decreased availability of suitable habitat, which may be more limited between these areas. Sleeping Bear Dunes and nearby surrounding areas make up a large contiguous amount of land that has been preserved by the National Parks Service, The Nature Conservancy, and other local land conservancies. Thus, the dune habitat is often continuous, with limited human development. On the other hand, Traverse City, Petoskey State Park, and Grand Marais areas have more human development along the lakeshore, which may provide additional barriers to gene flow among these populations.

Management histories could also be contributing to the differences in genetic diversity seen among these populations of baby’s breath. The entire Petoskey State Park population was treated with herbicide or manual removal from 2007 – 2012 by The Nature Conservancy. At this time, managers considered the population to be at a desirable management level, and it has been unmanaged since (TNC 2012 internal report). It is possible that the intensive management resulted in a population bottleneck, and the population rebound following 2012 came from a reduced number of individuals leading to the reduced genetic diversity that we observe today. However, this is probably not the only reason for the lower levels observed. The Arcadia Dunes and Zetterberg Preserve populations have also been regularly managed since 2004 and 2007, respectively, so if management is solely driving these patterns we would expect Arcadia Dunes and Zetterberg Preserve to also have reduced genetic diversity. Although the Arcadia Dunes population does have the lowest allelic richness and heterozygosity of all the populations in cluster 2 (Figure 2), both populations have relatively high genetic diversity despite over ten years of management. It is possible that higher levels of gene flow between these populations and those in Sleeping Bear Dunes may be helping to maintain genetic diversity. F_ST_ values between Zetterberg Preserve and other populations in Sleeping Bear Dunes range from 0.017 – 0.090, suggesting some gene flow, particularly with the population at the southern boundary (SB) of Sleeping Bear Dunes. Furthermore, infestations on private properties adjacent to Zetterberg Preserve have presumably buffered the population sizes. Given Petoskey State Park’s geographic distance from Sleeping Bear Dunes, limited gene flow between them would prevent the maintenance of high genetic diversity after intense management.

The topography and habitat heterogeneity of the dune system likely contributes to the pattern of population structure of baby’s breath throughout the Michigan dunes. Habitat heterogeneity can drive population structure, with variation in habitat type within the dunes acting as barriers to dispersal (Henry et al. 2009, Fant et al. 2014). Baby’s breath is typically found in open back dune habitat, but has also been found in the fore dunes close to the lake beach and on steep dune sides. However, forested areas that are part of the back dunes have been identified by land managers as barriers between populations, preventing population spread of baby’s breath (personal communication, Shaun Howard and Jon Throop). This can lead to populations in relatively close proximity to one another showing high levels of genetic differentiation and is likely a contributor to the significant population structure found among the populations in Sleeping Bear Dunes. For example, the Empire Bluff population (EB) is located on the tip of a dune bluff: a small visitor outlook point surrounded by forest, and seems to be isolated from nearby populations. Despite its geographic proximity to Platte Bay (PB) (8.22 km), it is more genetically similar to Sleeping Bear Point (SBP), a population 12.73 km away (F_ST_ = 0.106 and F_ST_ = 0.069 respectively).

A Mantel test for isolation by distance (IBD) revealed a moderate positive relationship between nSSR genetic distances (based on transformed pairwise F_ST_ values) and geographic distances (straight-line distances in km) of all populations (Figure 4a), and was also found for the analysis of populations in STRUCTURE cluster 2 (Figure 4c). However, when examining the IBD relationship within cluster 1 from the STRUCTURE analysis, this positive relationship is not significant (Figure 4b). We attribute the overall significant relationships found (Figure 4a and 4c) to the strong genetic differences between populations in the two main clusters, as well as the genetic difference of the Arcadia Dunes population when compared to Zetterberg Preserve and Sleeping Bear Dunes populations. Though geographic distance possibly influences the strong structuring of distant populations, the isolating effect of the topography within the dunes could have an effect that overrides that of geographic distance, particularly on smaller spatial scales such as that observed in Sleeping Bear Dunes. These results further support the strong regional differences between the two clusters identified in the Bayesian analysis (Figure 2).

The tumbleweed mechanism of dispersal that baby’s breath employs could be an effective means to disperse seeds, but it is possible that the strong topographical structure, habitat heterogeneity and variable weather patterns within the dunes impact seed dispersal for gene flow more than they impact pollination. Baby’s breath has been found to attract a diverse array of pollinator species (Baskett et al. 2011, Emery and Doran 2013), sometimes at the expense of native plant pollination, while seed dispersal is primarily limited to wind-driven tumbleweeds. The variation in **Φ**_ST_ values between the two marker types (nSSR **Φ**_ST_ = 0.355, cpSSR **Φ**_ST_ = 0.736) indicates that barriers to seed dispersal may be more limiting for gene flow than pollination. Darwent (1975) also suggested that though seeds could be dispersed up to 1 km, many of the seeds were released near the parent plant prior to the stems tumbling. This could result in strong population structure due to a lower frequency of migrants. Therefore, the elements of the dune ecosystem could be impacting gene flow through seed dispersal by further limiting the plant’s ability to spread throughout the landscape. However, these comparison of cpSSR results to nSSR should be taken with some caution, as we had a limited number of polymorphic cpSSR markers. Though we chose to use microsatellites within the chloroplast genome to increase the likelihood of polymorphism, we still found these regions to be well-conserved and with limited variation in our dataset. Therefore, we cannot rule out the possibility of fragment size homoplasy confounding results of low genetic diversity in some populations (Bang and Chung 2015).

### Structure analysis

Our results from the Bayesian clustering analysis in STRUCTURE (Pritchard et al. 2000, Evanno et al. 2005) separate the populations of baby’s breath along the Michigan coastal dunes into two genetic clusters (*K* = 2) (Figure 2). A similar pattern was found when the nSSR dataset was analyzed using a PCoA (Supplemental C) and a DAPC (Figure 3), with the exception that the individuals of the Arcadia Dunes population further separated from the Sleeping Bear Dunes and Zetterberg Preserve individuals in the latter analyses. The clusters are mainly divided into the Traverse City, Petoskey State Park, and Grand Marais cluster (cluster 1) and the Sleeping Bear Dunes populations, Zetterberg Preserve, and Arcadia Dunes cluster (cluster 2). The distribution of cpSSR haplotypes (Figure 5) across populations further illustrates the strong genetic clusters present in this dataset. Specifically, some haplotypes are only found in certain populations and within each main population cluster. Haplotypes 1, 2, and 4 only occur in populations in cluster 2. Haplotypes 3 and 5 only occur in cluster 1, with haplotype 3 being private to the five individuals in Grand Marais that were located separately from the rest of those sampled at this location (Figure 5). The two distinct population clusters are separated by the Leelanau peninsula, which may be helping to limit gene flow among these clusters. This partitioning of cpSSR haplotypes could be due to seed dispersal limitations from habitat fragmentation, unsuitable habitat, and land use, as the peninsula is comprised mainly of private residential properties along the narrow shoreline.

Understanding the invasion history of a species can help shed light on the factors and processes that contributed to the success of the species establishment. For baby’s breath, it has been assumed that invasive populations were the result of ornamental plants escaping from gardens or being purposely planted for horticultural means (personal communication with TNC managers). Whether the clusters we observed for our dataset are the result of multiple independent introductions or the result of one introduction followed by serial invasions is not known. Given that populations along coastal Michigan cluster into two distinct groups, either scenario is possible (Lombaert et al. 2018). In the serial invasion scenario, a founder population would have colonized one site in the Michigan coastal dunes, and then migrants from that population would have invaded subsequent areas (Lombaert et al. 2018). Over time, with limited gene flow, these populations could have become distinct and structured. However, we think this scenario may not be the best explanation for this invasion. Based upon herbarium records, the first occurrence of baby’s breath in northwest Michigan was recorded in 1913 in Emmet County where Petoskey State Park is located (Emmet Co., 1913, catalog: 355638, *Gleason s.n.*, MICH). Records from Leelanau and Benzie counties, where Sleeping Bear Dunes, Zetterberg Preserve and Arcadia Dunes are located, were not collected until the late 1940’s (Leelanau Co., 1947, catalog: 355348, *P.W. Thompson L-302,* MICH). If Petoskey State Park was the founding population for this invasion, we would expect higher genetic diversity in this population relative to those in Sleeping Bear Dunes, Zetterberg Preserve, and Arcadia Dunes, since a serial introduction would result in additional bottlenecks from the founding population. However, we observed the opposite pattern of genetic diversity. Additionally, there are private cpSSR haplotypes to each of these two clusters (Figure 5), a pattern we would not expect to see if all the populations came from one introduction event.

The other invasion scenario describes at least two independent introductions to the Michigan coastal dunes (Lombaert et al. 2018). In this scenario, we would expect strong genetic differentiation between the two or more founding populations. Our data supports this, as we observed both nSSR and cpSSR alleles privately shared only between populations within the same cluster. In addition, for the cpSSR markers, distinct haplotypes were found between the two regions, with haplotype 5 only observed in the Grand Marais, Petoskey State Park, and Traverse City cluster while haplotypes 1, 2, and 4 were only found in the Sleeping Bear Dunes, Zetterberg Preserve, and Arcadia Dunes cluster. There was also a high proportion of nSSR alleles common to both clusters, but this could be the result of limited genetic diversity in the initial source populations (Allendorf and Lundquist 2003). This scenario is particularly plausible, as the source populations would likely be a type of horticultural strain, given the popularity of perennial baby’s breath in the floral industry (Vettori et al. 2013, Calistri et al. 2014). This hypothesis of at least two independent introductions also agrees with the herbarium record: a potential introduction event could have occurred in the early 1910’s, leading to cluster 1 (GM, PS, TC), and a separate introduction event could have occurred in the late 1940’s, leading to the establishment of the populations in Zetterberg Preserve and Sleeping Bear Dunes (cluster 2).

In addition to supporting the identified patterns in the nSSR dataset produced from the STRUCTURE analysis, the PCoA and DAPC (Supplemental C and Figure 3) allowed us to identify more subtle population structuring. Specifically, the PCoA (Supplemental C) illustrates the Arcadia Dunes population separating from the other populations along principal coordinate 2. The DAPC (Figure 3) also shows the subtler variation among populations within the Sleeping Bear Dunes populations (specifically Figure 3c), and continues to support the segregation of the Grand Marais, Petoskey State Park, and Traverse City populations from the rest that we see in the STRUCTURE analysis (Figure 2). Variation in allele frequencies and decreased allelic richness are two factors that could explain the divergence of the Arcadia Dunes population in the PCoA (Supplemental C); there are no private alleles or other obvious patterns causing this population to cluster separately from nearby populations (Zetterberg Preserve and South Boundary in Sleeping Bear Dunes). The higher rates of admixture between the two main clusters in Arcadia Dunes individuals (Figure 2) could also be a reason for its slight divergence from cluster 2. However, what is driving this potential higher level of admixture in the Arcadia Dunes population compared to others is currently unknown. Arcadia Dunes is a popular recreation area among locals and tourists (personal communication Jon Throop, Grand Traverse Regional Land Conservancy). Additionally, the autumn season brings about a high volume of foot traffic through all the dune areas of Michigan. It is possible that people may be accidentally transporting baby’s breath seeds between these otherwise isolated populations, as the seed phenology coincides with the autumn senescence. While human transport of seeds may be occurring at other locations as well, Arcadia Dunes is a small enough population that newly introduced genotypes could have a higher likelihood of being detected from sampling relative to other larger populations, such as one in Sleeping Bear Dunes.

The invasion of baby’s breath to the Great Lakes has the potential to disrupt the dynamism of the dune landscape and biological community in northwest Michigan, and this threat has led to increased concern over its pervasiveness regionally and nationally. Estimating the genetic structure of invasive populations can lead to a better understanding of the invasion history and the factors influencing the success of an invasion (Crosby et al. 2014, Piya et al. 2014, Sakata et al. 2015). Through population level analysis, we found strong genetic structure present that separates the invasion in the Michigan dunes into two main regions. Based on these results, we suggest that the contemporary baby’s breath population within the Michigan coastal dune system is the result of at least two separate introduction events. The genetic structure identified for these baby’s breath populations probably results from a combination of demographic processes –multiple introductions, bottleneck events, isolation, and admixture, along with landscape level processes. The topography of the dunes is heterogeneous but also constantly shifting, and the baby’s breath invasion is one example of how this dynamic system can shape the establishment, gene flow, and spread of invasive plant populations.

## ACKNOWLEDGEMENTS

The authors thank the Environmental Protection Agency – Great Lakes Restoration Initiative Grant (C.G.P., Grant #00E01934) for financial support. We thank Dr. James McNair for assistance with statistical analyses, Dr. Timothy Evans for assistance with herbarium data expertise and submission, and Kurt Thompson for assistance with geographic data collection and visualization. Finally, we thank Emma Rice for all her assistance in field data collection and Matthew Kienitz for his assistance in field and laboratory data collection.

## DATA ARCHIVING

All genotype data will be submitted to the Dryad Digital Repository upon acceptance. Nuclear microsatellite sequences have been deposited to GenBank (Table 1).

---

**Supplemental A.**
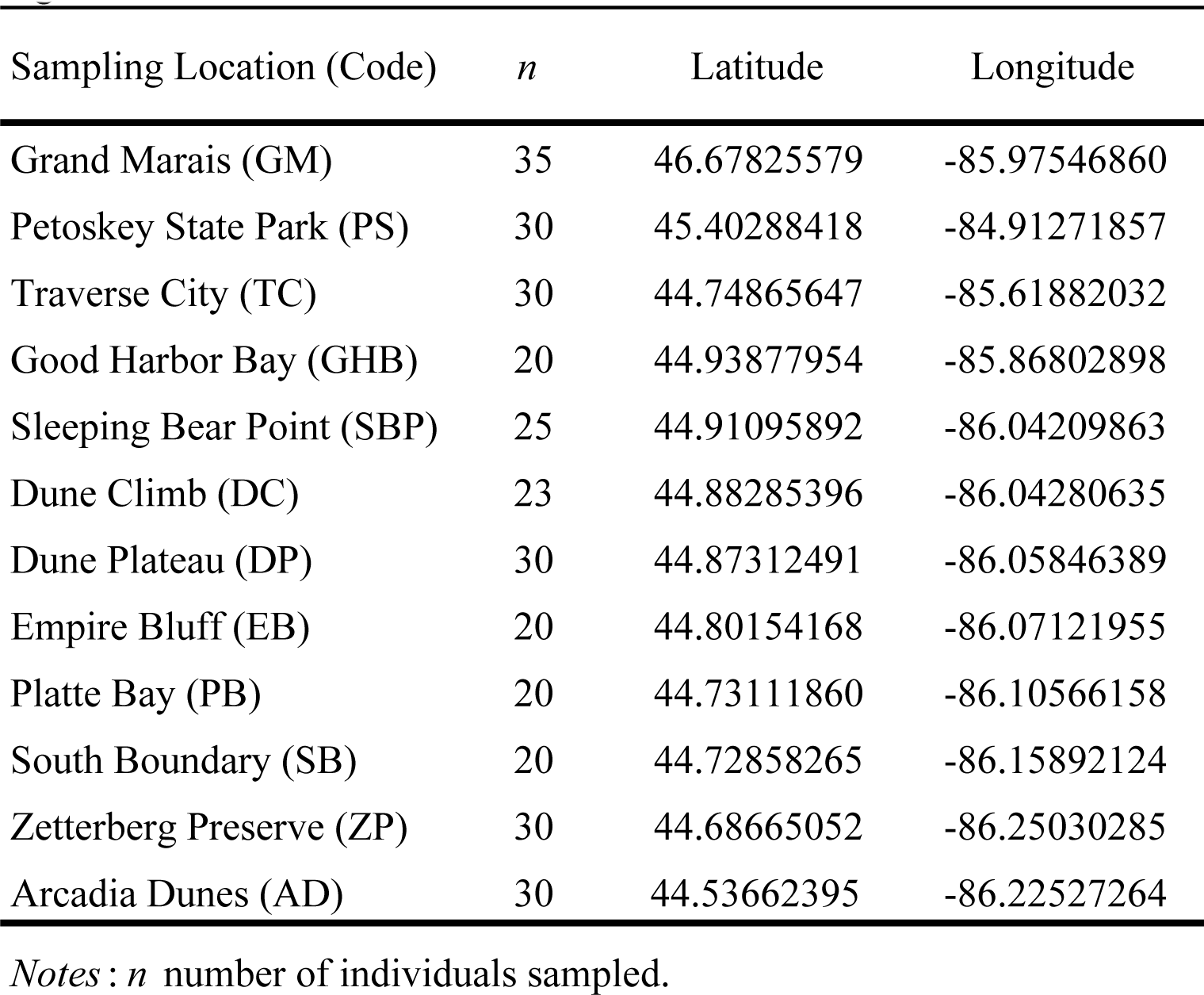
Sampling location names and geographic coordinates for baby’s breath analyzed in this study. All locations are in Michigan. Location abbreviations are used in the main text and following tables and figures.

**Supplemental B.**
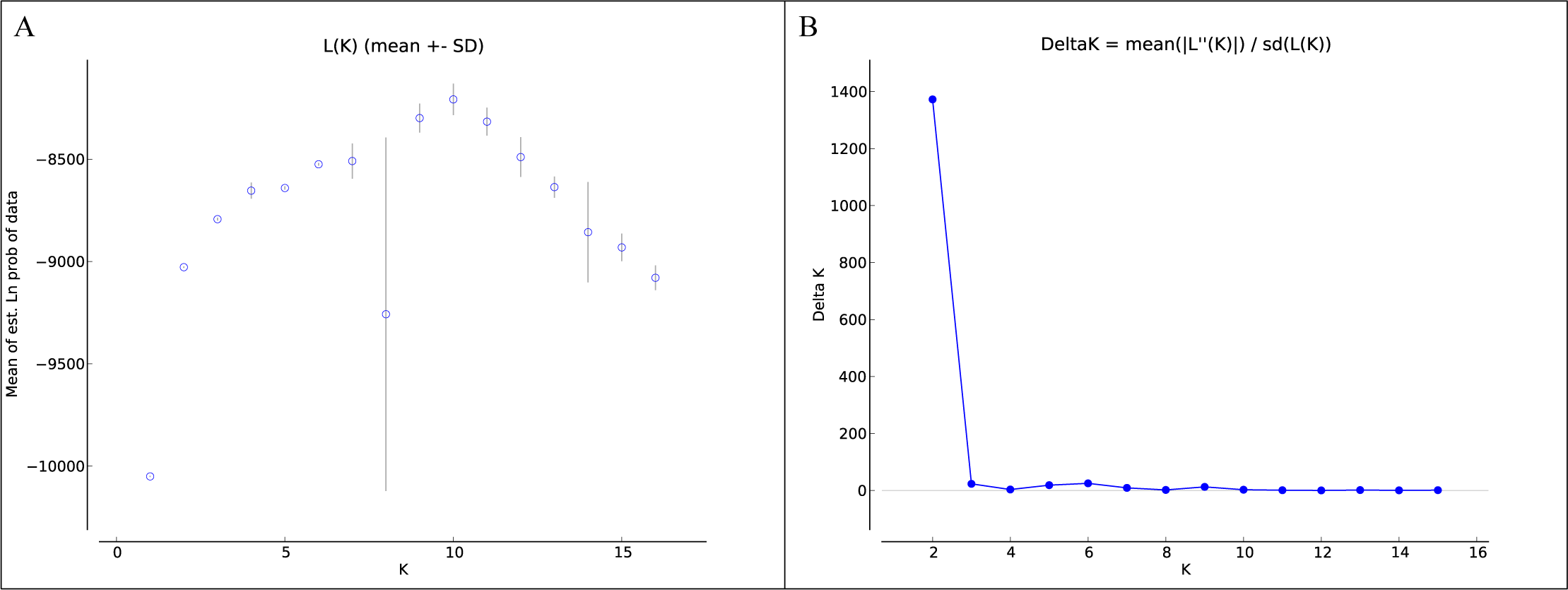
Bayesian clustering analysis of all 12 baby’s breath populations from the program STRUCTURE (Pritchard et al. 2000). (A) Mean L(K) (± SD) over 10 runs for each value of *K.* (B) Plot of Evanno’s ΔK method (Evanno et al. 2005) where the largest rate of change suggests the highest likelihood of cluster number. This analysis was run without inferring any prior information on sampling location, and two genetic clusters were inferred from this data.

**Supplemental C.**
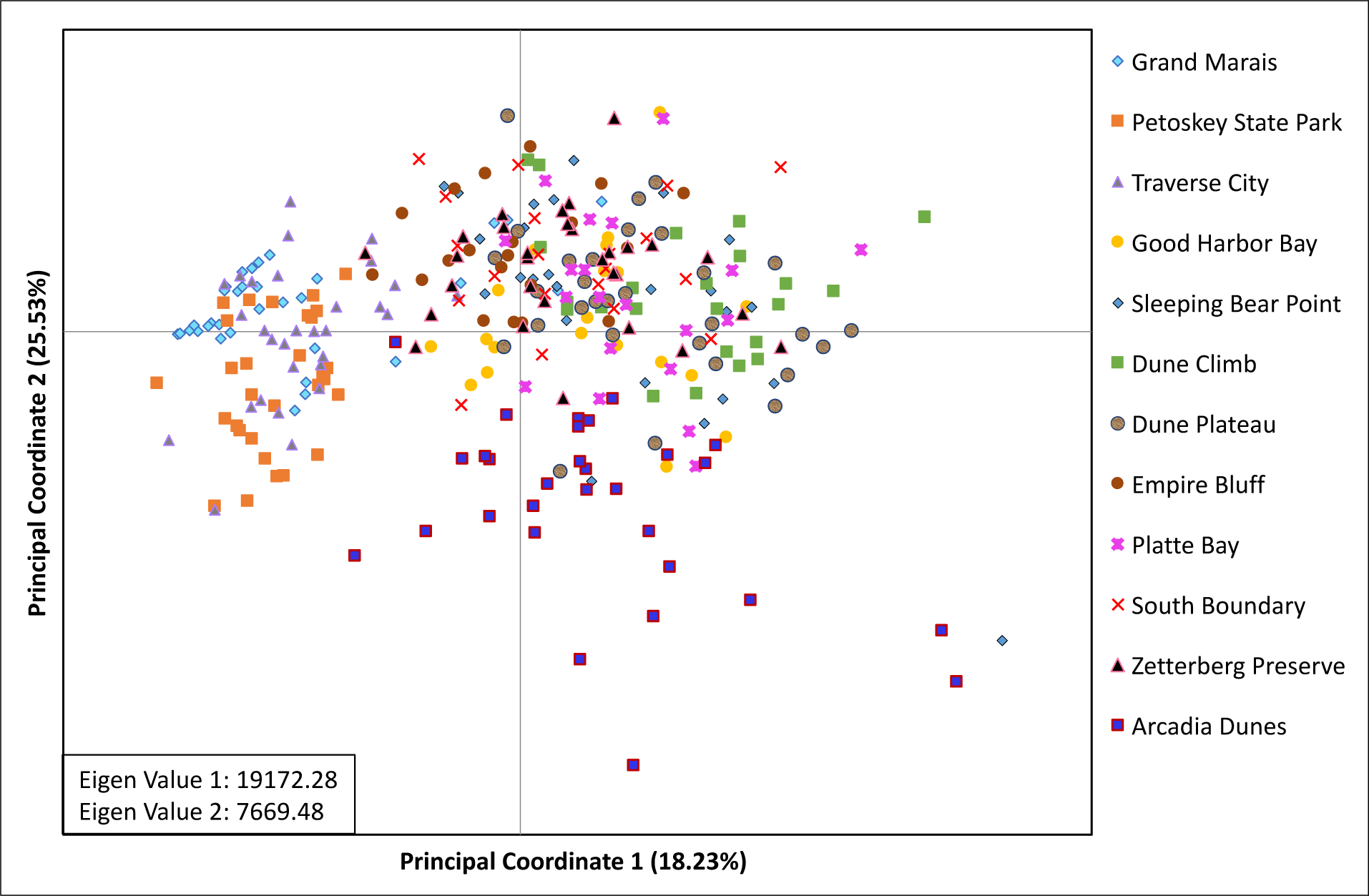
Principal Coordinates Analysis (PCoA) based on a genotypic distance matrix between all baby’s breath individuals performed in GenAlEx 6.502 (Peakall and Smouse 2006, 2012). Individuals labeled by sampling location.

**Supplemental D.** Analysis of Molecular Variance (AMOVA) for 14 nuclear and 2 chloroplast SSR loci in 12 populations of baby’s breath. Regional differences identified in the Bayesian clustering analysis (*K* = 2) were included in AMOVA, and the analysis was based on ! estimates to compare variance across both marker types following Excoffier et al. (1992) and Weir and Cockerham (1984).

| Source of Variation | nSSRs |  |  |  | cpSSRs |  |  |  |
| --- | --- | --- | --- | --- | --- | --- | --- | --- |
| | df | % of Variation | $\Phi$ – statistics | P – value | df | % of Variation | $\Phi$ – statistics | P – value |
| Among regions | 1 | 22.62 | 0.226 | 0.0001 | 1 | 26.27 | 0.263 | 0.0001 |
| Among populations within regions | 10 | 12.89 | 0.167 | 0.0001 | 10 | 47.37 | 0.643 | 0.0001 |
| Within populations | 301 | 64.49 | 0.355 | 0.0001 | 301 | 26.36 | 0.736 | 0.0001 |

